# Elevated surface La promotes hyperfusion and contributes to impaired resorption in osteopetrosis

**DOI:** 10.1101/2025.09.07.674639

**Authors:** Jarred M. Whitlock, Evgenia Leikina, Hong-Yin Wang, Wendy Zhang, Griffin Katz, Nina Reuven, Ari Elson, Benjamin Geiger, Leonid V. Chernomordik

## Abstract

The skeleton is a living, biological tissue responding to the biomechanical demands placed upon it throughout life. The individual bones creating this physiological system are each shaped by a multinucleated cell type - the osteoclast - that sculpts each bone in collaboration with local cellular partners, which offer chemical and even tactile feedback of many sorts. Unfortunately, the perturbation of osteoclast formation and function underpins a broad range of human skeletal pathologies, including osteopetrosis - a systemic pathology characterized by impaired osteoclast resorption leading to skeletal thickening, brittle bones, frailty, and lethality. Here, we describe a molecular dysfunction observed in murine and human models of two forms of osteoclast-rich, autosomal recessive osteopetrosis, and our approach for exploiting this molecular dysfunction to correct pathologic osteoclast hyperfusion and resorptive impairment. We find that La - a manager of osteoclast fusion and subsequent resorptive activity - is greatly elevated at the surface of osteoclasts upon loss of *SNX10* or *OSTM1*. Using inhibitory antibodies, we suppress excessive La surface function in these mutant osteoclasts, impede osteopetrotic hyperfusion and restore osteoclast resorptive function. We share these observations as proofs-of-principle that osteoclast fusion represents a viable therapeutic target for addressing osteoclast dysfunction in diseases underpinned by excessive osteoclast multinucleation and perturbed resorptive function.

## Introduction

Lifelong bone remodeling, which is essential for maintaining physiological skeletal functions is accomplished by a tightly-regulated interplay between multinucleated osteoclasts (OCs) that resorb old, frail bone and mononucleated osteoblasts that deposit new bone^1^. OC-osteoblast collaboration is required for postnatal skeletal development and completely regenerates most of the skeleton every decade of life to maintain its biomechanical integrity and function^2^. OCs largely set the rate of skeletal remodeling^3^, their resorptive activity is essential for vertebrate life^4-6^, and their malformation and dysfunction commonly leads to skeletal disease. Specifically, excessive OC resorptive activity leads to pathologically low bone mass (e.g., in osteoporosis) or reduced resorptive activity leads to the formation of highly dense bone (e.g., in osteopetrosis).

OCs arise from circulating monocytes that differentiate into macrophage precursors^7, 8^. Precursors enter osteoclastogenesis following activation by receptor activator of nuclear factor-kappa B ligand (RANKL)^7^. During osteoclastogenesis, committed OCs fuse plasma membranes (PMs) and produce multinucleated syncytia with 5-12 nuclei/cell (i.e., a process we refer to here as multinucleation)^9, 10^. The number of fusion events forming an OC determines both its nuclear number and, crucially, its capacity to resorb bone^10-12^. Importantly, many skeletal diseases are underpinned by perturbations in the number/size of OCs that drive pathological skeletal remodeling and lead to pain, dysfunction, and even death^10, 13-18^.

Notably, there is growing information reported about cell fusion during osteoclastogenesis^19, 20^, as well as ample mechanistic data concerning the basic mechanisms underlying the bone resorption process^1, 21^, yet the temporal interplay and functional interdependence between these processes remains largely obscure. Better refinement in the fundamental understanding of processes regulating multinucleation is urgently needed to resolve how OC number, size, and activity is controlled. This resolution will lead to the identification and develop next generation targets for correcting pathologic skeletal remodeling in disease while maintaining healthy OCs and their vital contribution to life-long skeletal development, adaptation, and recovery.

We discovered that OCs repurpose a nuclear RNA chaperone - Lupus La protein (La) - as a fusion manager at their surface, setting OC size and activity^22^. Canonically, La - the *SSB* gene product - is an abundant, essential, nuclear RNA-binding protein that facilitates the maturation of non-coding RNA polymerase III transcripts^23^. Recently, we discovered an “*<u>unconventional” function where La promotes OC multinucleation and bone resorption</u>*. In La’s unconventional OC role, its C-terminal nuclear localization sequence (NLS) is cleaved away in response to RANKL-activated osteoclastogenesis. This cleaved La traffics to the OC surface and promotes cell-cell fusion, determining OC nuclear number and resorptive activity. When OCs reach a mature size, La is removed from the cell surface, new, intact La traffics to the nuclei of mature OCs via its NLS, and fusion between mature OCs is suppressed^22^.

In contrast to bone loss diseases, pathologic shifts in the balance between bone-forming and bone-resorbing activities in osteopetrosis lead to insufficient resorption, abnormal growth and overly dense bones^24^. In diverse types of hereditary osteopetrosis, increased bone density is caused by abnormalities in OC formation and function. Intriguingly, the paradigm suggesting higher bone resorption activity for larger OCs is broken by recent reports that some types of osteopetrosis are accompanied by the formation of giant, inactive osteoclasts^18, 25, 26^. OC-rich autosomal recessive osteopetrosis (OC-rich ARO), a genetic disease in which OCs are formed but cannot resorb bone^24^, can be caused by mutations in, among other proteins, the vesicular-trafficking protein sorting nexin 10 (SNX10)^27, 28^ or the osteopetrosis associated trans-membrane protein 1 coding sequence (*OSTM1*)^24, 29^. Loss of bone-resorbing activity in both *SNX10*- and *OSTM1*-linked OC-rich ARO is accompanied by hyperfusion, leading to a hundreds-fold increase in the sizes of osteoclasts in culture^30, 31^; large OCLs are also detected in vivo in mice lacking SNX10^30^. Yet, despite their gigantic size, these OCs cannot degrade bone^32,33^. OC-rich ARO is a lethal pediatric disease, with most patients passing in the first decade of life. While bone marrow transplantation is an effective treatment for some sub-types^24, 34^, OC-rich ARO has no currently approved therapeutic, and many patients are provided little more than palliative care. Current efforts towards the development of stem cell based therapeutic interventions show promise for future interventions^34^, however, ARO represents a lethal pediatric skeletal disease with urgent, unmet therapeutic needs.

Typically, the loss of OC resorptive capacity in OC-rich ARO is discussed as a consequence of impaired function or delivery of acidification machinery to the OC membrane ^24, 34^. Recently, OCs in several forms of OC-rich ARO were shown to exhibit an “immature” behavior, constantly fusing and never forming resorption pits^31, 33^. A carefully regulated multinucleation mechanism is tasked with producing OCs of an appropriate size to match the biomechanical needs of the remodeling skeleton. Previously, Barnea-Zohar et al. suggested that OC fusion stops when OC fusion machinery is removed from the cell surface, possibly by endocytosis, and that *SNX10* deficiency promotes fusion by reducing endocytic activity^31^. OSTM1 also plays a putative role in intracellular protein trafficking^35^. In addition, non-osteopetrosis linked genes have been suggested to regulate OC fusion via endocytosis (e.g., CD13). The role of CD13 in regulating endocytosis has also been demonstrated where OC multinucleation is halted by the internalization of the fusion related protein DC-STAMP and loss of CD13 promotes OC multinucleation^36^. We propose that these findings indicate that hyperfusion is not a parallel phenotype but directly contributes to the loss of OC resorption and skeletal malformation in OC-rich ARO.

In this study, we explore how *SNX10* and *OSTM1* deficiencies dramatically promote multinucleation in murine and human osteoclasts. We specifically tested the hypothesis that loss of SNX10 or OSTM1 promotes the retention of La at the surface of mature mutant OCs, thereby inducing a sustained hyperfusion phenotype that eventually leads to impaired resorptive function. We found in immortalized or primary murine and human systems that loss of *SNX10* or *OSTM1* leads to the elevation of La at the surface of osteopetrotic OCs. In each case, inhibition of surface La via inhibitory monoclonal antibodies rescues the OC hyperfusion phenotype associated with OC-rich ARO. Moreover, we demonstrate that suppressing excessive multinucleation via La inhibitory antibodies or peptides that inhibit syncytin 1-an endogenous retroviral protein previously recognized for its role in OC fusion^37, 38^ - not only rescues osteopetrotic hyperfusion but also improves resorptive function. These findings demonstrate that loss of SNX10 or OSTM1 leads to La retention at the surface of OCs, which promotes hyperfusion. Moreover, our data suggest that hyperfusion directly contributes to the loss of OC resorption and skeletal malformation in OC-rich ARO and indicate that fusion represents a therapeutic target with significant promise for treating OC-rich ARO.

## RESULTS

### Suppression of Snx10 leads to osteoclast hyperfusion and elevated La protein

To evaluate how the loss of *Snx10* impacted the OC multinucleation manager La protein, we suppressed *Snx10* via siRNA-mediated knockdown in Raw 264.7 cells during RANKL activated osteoclastogenesis. RNA interference decreased the steady-state level of Snx10 (Fig. 1a) and, as described earlier^31^, dramatically promoted fusion and multinucleation in OCs (Fig. 1b,d). Intriguingly, *Snx10* knockdown led to a concomitant increase in the steady-state level of La (Fig. 1a). To assess where this elevated La protein localized in Raw 264.7 during osteoclastogenesis, we performed immunostaining with antibodies raised against La (α-La). Under permeabilizing conditions, we observed no overt change in La cytosolic staining. In stark contrast, under non-permeabilizing conditions we found that SNX10 suppression increased La staining at the surface of Raw 264.7-derived OCs (Fig. 1c). To assess whether elevated levels of La contributed to the osteopetrotic hyperfusion observed upon *Snx10* suppression, we simultaneously suppressed both *Snx10* and *Ssb* (La transcript). We found that simultaneous suppression of *Ssb* partially rescued the aberrant OC multinucleation caused by *Snx10* knockdown (Fig. 1d). Taken together, these data suggest that the hyperfusion associated with loss of *Snx10* is accompanied by an increase in La at the surface of OCs, which promotes this hyperfusion.

**Figure 1:**
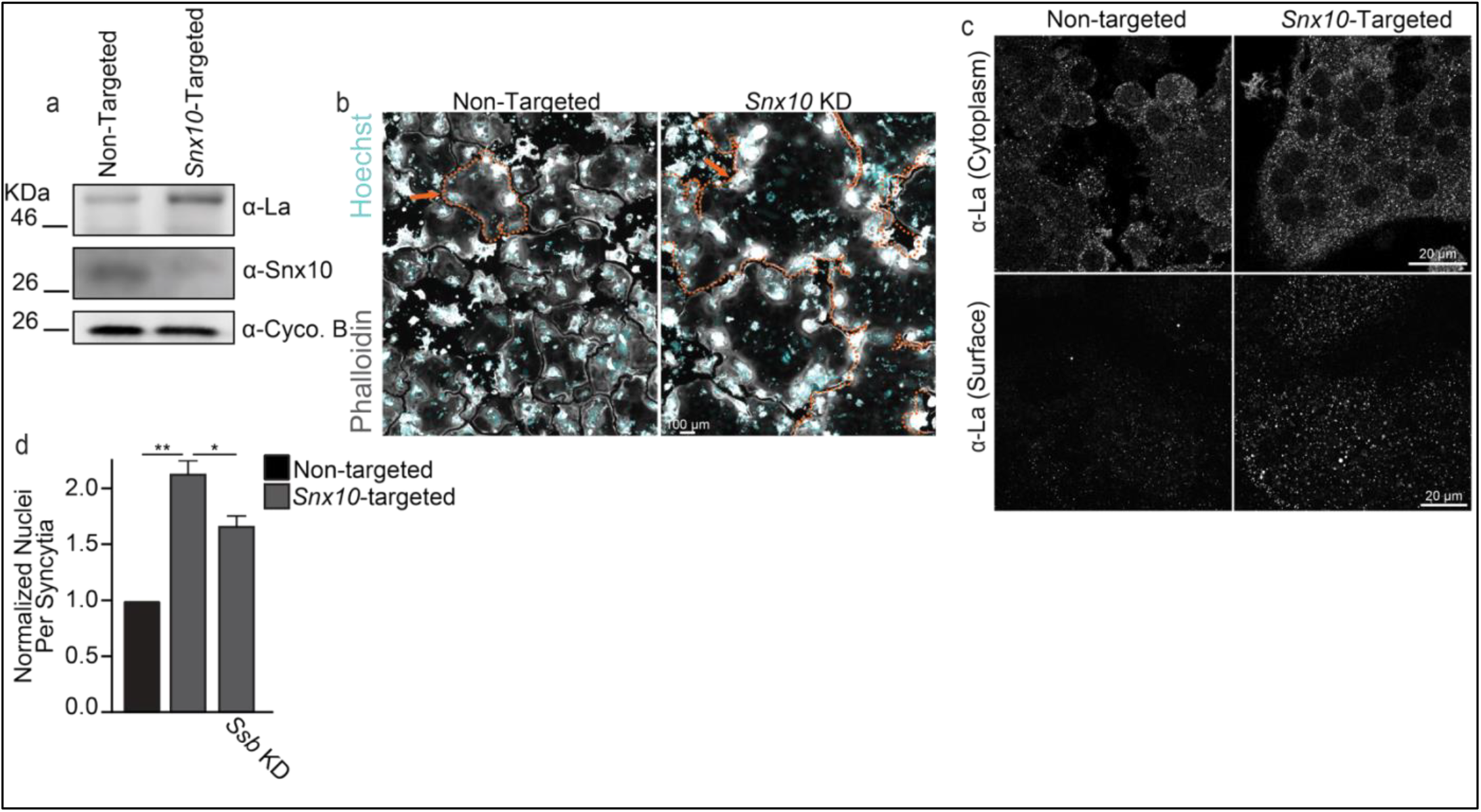
*Snx10* loss elevates surface La and promotes osteoclast hyperfusion. **(a)** Representative Western blot depicting differences in the steady-state levels of La, Snx10, and the loading control Cycophilin B. **(b)** Representative immunofluorescence images depicting α-La staining intensity in permeabilized (top) vs non-permeabilized (bottom) conditions in osteoclasts treated as in **a. (c)** Representative fluorescence images depicting the formation of multinucleated osteoclasts from RAW 264.7 cells treated as in **a. (d)** Quantification of the number of nuclei per syncytia in RAW 264.7 derived osteoclasts following the introduction of non-targeted, *Snx10*-targeted, or *Snx10*- & *Ssb*-targeted siRNA. n = 3. p = 0.002 & 0.03, respectively. Error bars = SEM

### Elevated surface La facilitates hyperfusion associated with Snx10 and Ostm1 loss

To extend our initial observation that the hyperfusion associated with *Snx10* loss was accompanied and promoted by elevation of La at the surface of OCs, we turned to knockout Raw 264.7 lines lacking either *Snx10* or *Ostm1* (described and characterized previously^33^). We characterized these knockout lines and confirmed that each exhibited loss of the respective *Snx10* or *Ostm1* gene products (Fig. 2a,b). Consistent with our *Snx10* knockdown results, we found that loss of either *Snx10* or *Ostm1* led to an increase in the steady-state level of La protein despite no similar increase in La transcript or the transcript levels of annexin A5 - a protein that anchors La to the surface of the cells^19^ (Fig. 2c & Supp. Fig. 1). This elevation in the steady state-level of La was again associated with a significant increase in the level of La staining at the surface of OCs upon *Snx10* loss (Fig. 2d,e). We observed a similar trend upon loss of *Ostm1*, however these massive osteopetrotic osteoclasts appeared unhealthy following hyperfusion and staining was variable (Fig. 2f, p = 0.087). Removal of one experiment where this trend was not observed (red dots) resulted in this trend reaching the level of significance (Fig. 2f removing red dots, p = 0.045). While the differentiation and fusion capacity of these Raw 264.7 lines exhibited variation from experiment to experiment, we consistently noted dramatic hyperfusion phenotypes in both the *Snx10* or *Ostm1* knockout lines relative to parental controls (Fig. 2g-j). Interestingly, the *Ostm1* knockout - modeling the most severe genetic linkage of OC-rich ARO - exhibited consistently elevated, faster OC hyperfusion relative to *Snx10* knockout (Fig. 2g,h vs i,j). However, in either case, application of α-La antibodies robustly suppressed hyperfusion and rescued OC size (Fig. 2 g-j). These findings further support the conclusion that elevated surface La, stemming from *Snx10* or *Ostm1* loss, is elevated on the surface of osteopetrotic OCs and facilitates hyperfusion.

**Figure 2:**
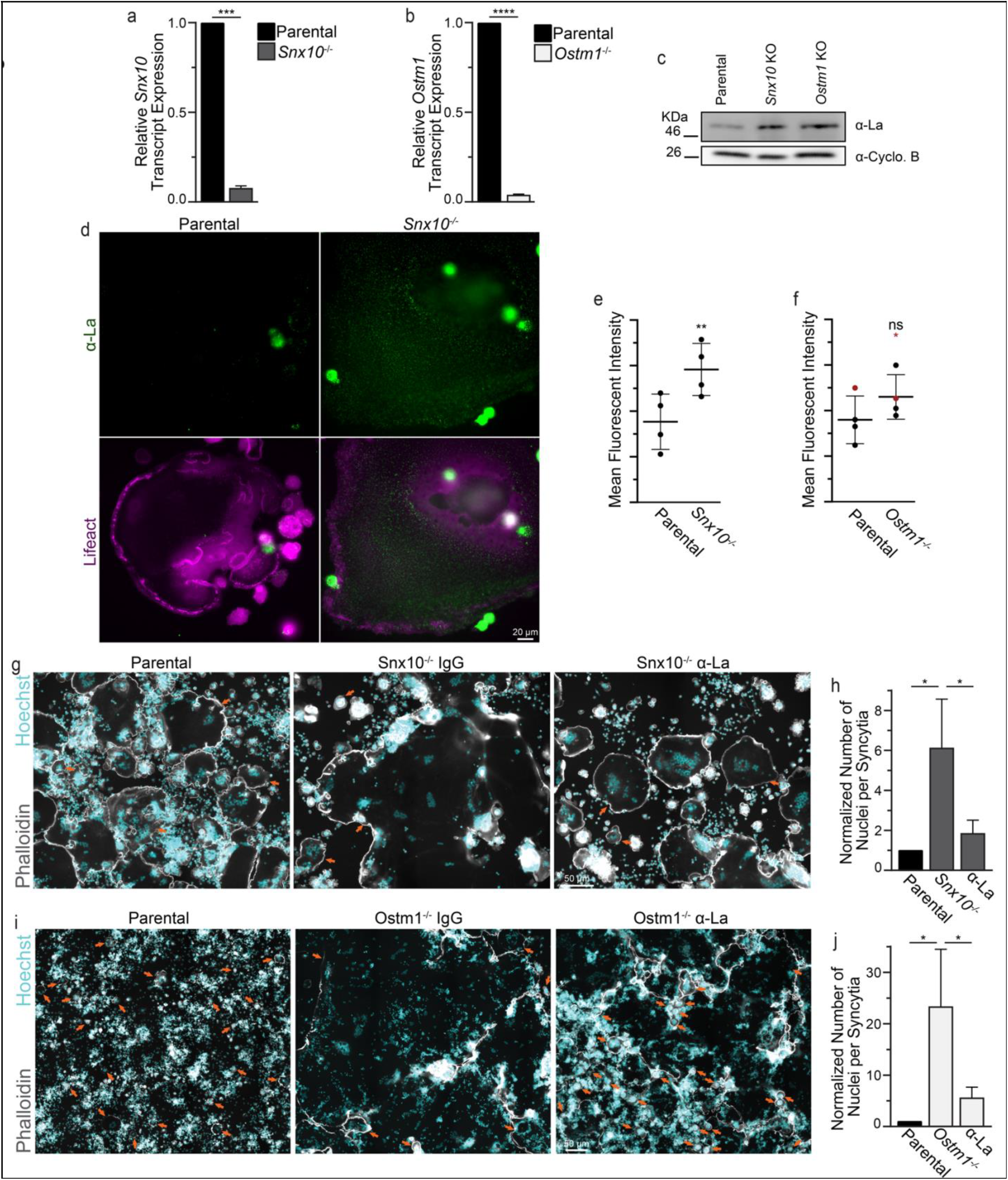
Elevated surface La facilitates osteoclast hyperfusion in response to deletion of *Snx10* or *Ostm1*. **(a)** Quantification of the steady-state level of *Snx10* transcript relative to *GAPDH*. n = 4. p = 0.0002 **(b)** Quantification of the steady-state level of *Ostm1* transcript relative to *GAPDH*. n = 5. p < 0.0001 **(c)** Representative Western blot depicting differences in the steady-state levels of La and the loading control Cycophilin B in lysates of RAW 264.7 derived osteoclasts. **(d)** Representative immunofluorescence images depicting α-La surface staining intensity in non-permeabilized conditions in parental vs *Snx10*^*-/-*^ osteoclasts. **(e)** Quantification of the average fluorescence intensity from **d**. n = 4. p = 0.007. **(f)** Quantification of the average fluorescence intensity of α-La surface staining in parental vs *OSTM1*^*-/-*^ osteoclasts. With, n = 4. p = 0.087. vs without red dots, n = 3. p =0.045. **(g)** Representative fluorescence images depicting the formation of multinucleated osteoclasts from RAW 264.7 cells with the application of α-La antibody vs isotype control (7.5μg/ml) **(h)** Quantification of the number of nuclei per syncytium normalized to parental control from **g**. (n = 4) (p = 0.01 and 0.02, respectively). **(i)** Representative fluorescence images depicting the formation of multinucleated osteoclasts from RAW 264.7 with the application of α-La antibody vs isotype control (7.5μg/ml). **(j)** Quantification of the number of nuclei per syncytium normalized to parental control from **i**. (n = 4) (p = 0.009 and 0.015, respectively). Error bars = SEM. Arrows denote individual OCs.

### Loss of SNX10 or OSTM1 elevates surface La and facilitates hyperfusion in human OCss

Next, we assessed the impact of suppressing *SNX10* in primary human OCs. Human osteoclasts were derived as previously described ^39^. Following RANKL activation, we introduced either non-targeting or *SNX10*-targeting siRNA and evaluated the impact of *SNX10* loss on La protein levels and human OC formation via fluorescence microscopy. Under permeabilizing conditions, we found that suppressing *SNX10* did not affect the cytoplasmic pool of La protein (Fig. 3a). In contrast, we found that, under non-permeabilizing conditions, *SNX10* loss significantly elevated the pool of La protein on the surface of human OCs. Concomitantly, we observed a shift in the formation of primary OCs, where the number of OCs formed were consistently reduced by ∼1/2 while the number of nuclei per OC nearly doubled (Fig. 3c-d). Similar to our immortalized, murine models, suppression of surface La activity via α-La antibodies suppressed hyperfusion and OC size under *SNX10* knockdown conditions (Fig. 3e,f). We also observed a dramatic elevation of the pool of La protein on the surface of human OCs following *OSTM1*-targeted siRNA delivery (Fig. 4a,b). Similar to our murine models, loss of *OSTM1* led to a more dramatic hyperfusion phenotype compared to *SNX10 (Fig. 4c,d vs 3e,f)*. For both *SNX10*-suppressed and *OSTM1*-suppressed OCs, α-La antibodies decreased the number of nuclei per syncytium, achieving nuclear numbers closer to those observed in control cells *(Fig. 4c,d vs 3e,f)*. However, while we consistently observed hyperfusion following *OSTM1* suppression, the variability in extent between replicates was such that these changes did not reach the level of significance (Fig. 4c,d). These data suggest that in human OCs, as in murine, surface La pools are elevated and contribute to the hyperfusion observed in response to *SNX10* and *OSTM1* loss.

**Figure 3:**
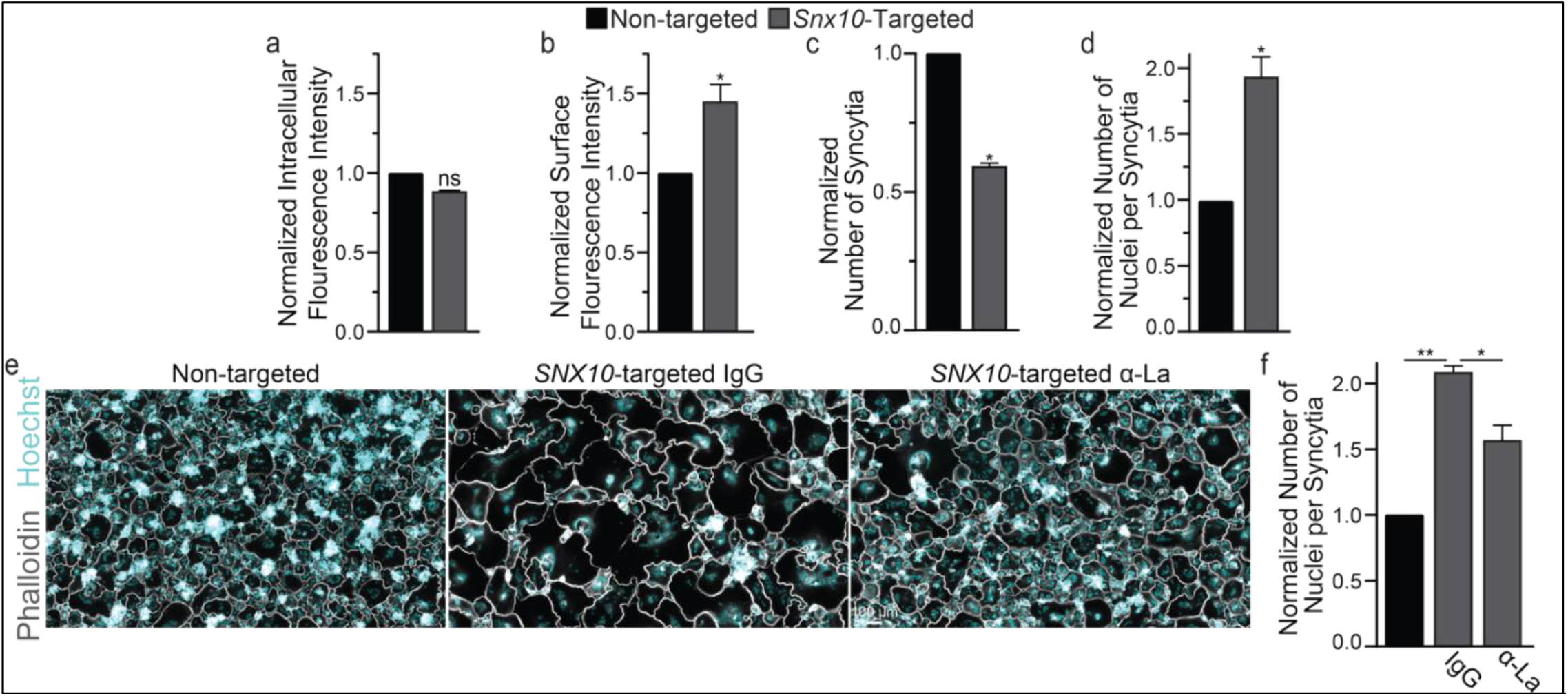
Suppression of *SNX10* elevates surface La and facilitates hyperfusion in human OC formation. **(a)** Quantification of α-La intracellular staining intensity in permeabilized osteoclasts. n = 2. p = 0.08 **(b)** Quantification of α-La surface staining intensity from non-permeabilized osteoclasts. n = 3. p = 0.02 **(c)** Quantification of the number of osteoclast syncytia. n = 3. p = 0.007 **(d)** Quantification of the number of nuclei per osteoclast syncytium from **c**. n = 3. p = 0.0008. **(e)** Representative fluorescence images depicting the formation of osteoclast syncytia with the application of α-La antibody vs isotype control. **(f)** Quantification of the number of nuclei per syncytium from **e**. n = 3. p = 0.002 & 0.013, respectively. (Normalized data = normalized to non-targeted control) (Syncytia = OC with 3+ nuclei) (antibodies = 7.5μg/ml) Error bars = SEM

**Figure 4:**
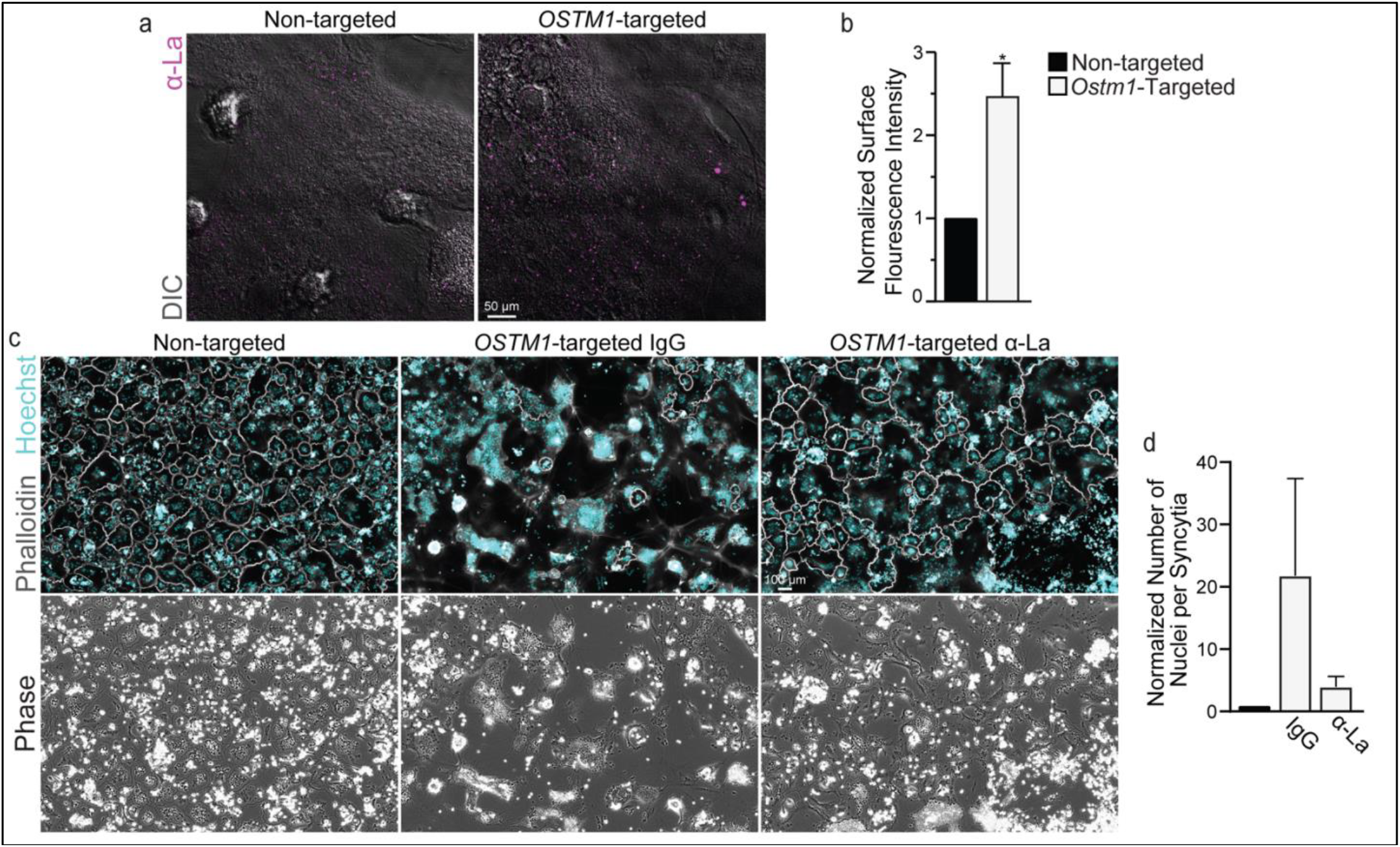
Suppression of *OSTM1* elevates surface La and facilitates hyperfusion in human osteoclast formation. **(a)** Representative immunofluorescence images depicting α-La surface staining intensity of non-permeabilized osteoclasts following siRNA application. **(b)** Quantification of fluorescent intensity from **a**. n = 3. p = 0.016. **(c)** Representative fluorescence images depicting the formation of multinucleated osteoclasts with the application of α-La antibody vs isotype control. **(d)** Quantification of the number of nuclei per syncytial osteoclast from **c**. n = 4 (Normalized data = normalized to non-targeted control) (Syncytia = OC with 3+ nuclei) (antibodies = 7.5μg/ml) Error bars = SEM

Previous hypotheses concerning the mechanistic underpinning of OC-rich ARO have focused on demonstrated and/or suspected perturbations in the trafficking of the machinery required for resorption pit acidification in osteopetrotic OCs rather than hyperfusion. While we suspect that alterations in acidification machinery (e.g., CLC transporters) contribute to the significant loss of resorptive remodeling in osteopetrotic bones, we hypothesized that hyperfusion also contributes to the loss of resorptive function in *SNX10*- and *OSTM1*-linked osteopetrosis. To test this hypothesis, we plated monocyte derived, human OCs on calcium-phosphate ceramics and introduced siRNA to model *SNX10* or *OSTM1* loss. Note that, in this *in vitro* model we only observed hyperfusion ∼4.5 days after RANKL application (∼2.5 days after siRNA introduction) and can only reliably measure osteopetrotic changes in mineral resorption at ∼5-6 day after RANKL application, leaving us a limited observational window for treatments. Because OCs begin forming at ∼2.5 days after RANKL application, OCs resorb mineral for 2x longer before the hyperfusion phenotype develops vs after. Despite this technical limitation, we find that suppression of *SNX10* or *OSTM1* results in ∼30% loss of mineral resorption (Fig. 5a,b). In both cases, suppressing hyperfusion by applying α-La antibodies rescued mineral resorption. Again, changes following suppression of *OSTM1* were reliably observed but variable in extent. Moreover, we also rescued resorption by applying a fusion inhibiting peptide^37, 38^ that blocks the function of another OC fusion protein - syncytin 1, suggesting that rescuing OC resorption is not reliant on specifically targeting La protein but rather suppressing osteopetrotic hyperfusion (Fig. 5c).

**Figure 5:**
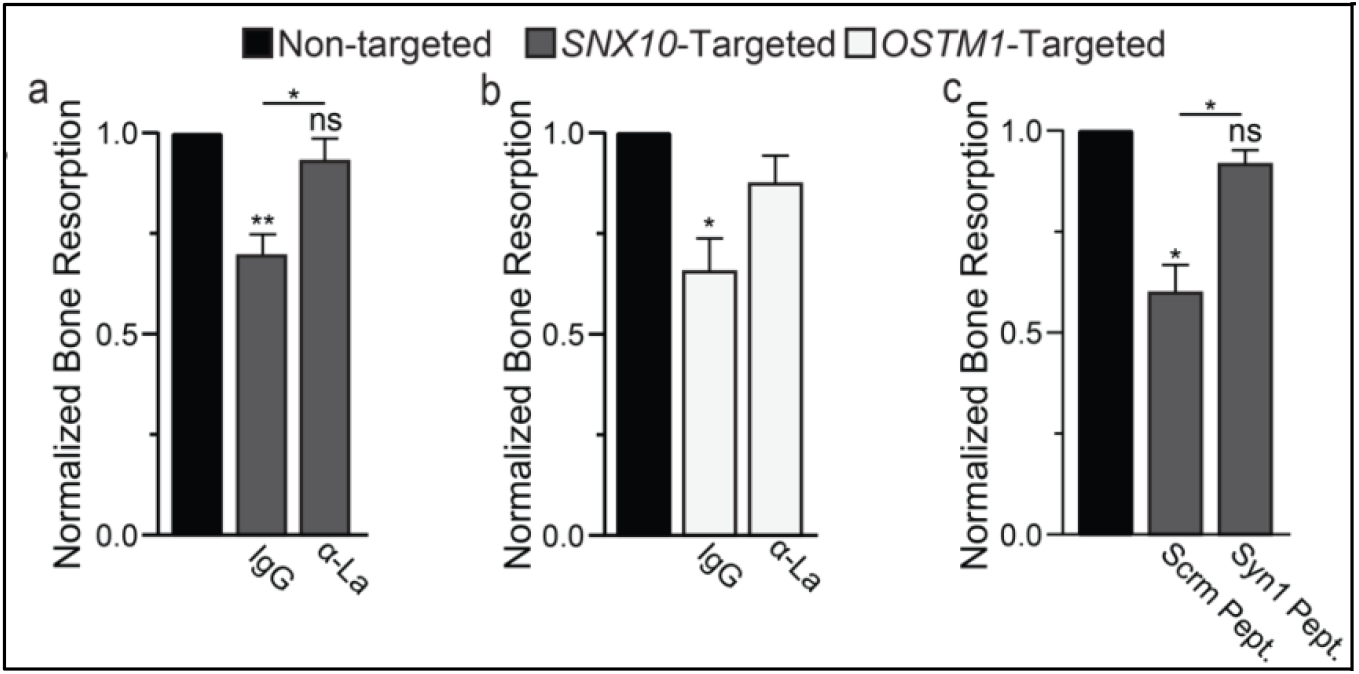
Suppressing hyperfusion rescues resorption activity of human osteoclasts stemming from *SNX10* and *OSTM1* loss. **(a)** Quantification of *in vitro* resorptive function of osteoclasts treated with either non-targeting or SNX10 targeting siRNAs and then incubated with either α-La or control IgG. n = 5. p = 0.003, 0.17 & 0.01, respectively. **(b)** Quantification of *in vitro* resorptive function of osteoclasts treated with either non-targeting or OSTM1-targeting siRNAs and then incubated with either α-La or control IgG. n = 4. p = 0.04 **(c)** Quantification of *in vitro* resorptive function of osteoclasts treated with either non-targeting or SNX10-targeting siRNAs and then incubated with either Syncytin1 inhibiting peptide (Syn1 Pept) or its scrambled version (Scrm Pept.). n = 3. p = 0.018, 0.22 & 0.03, respectively. (Normalized data = normalized to non-targeted control) Error bars = SEM.

**Figure 6:**
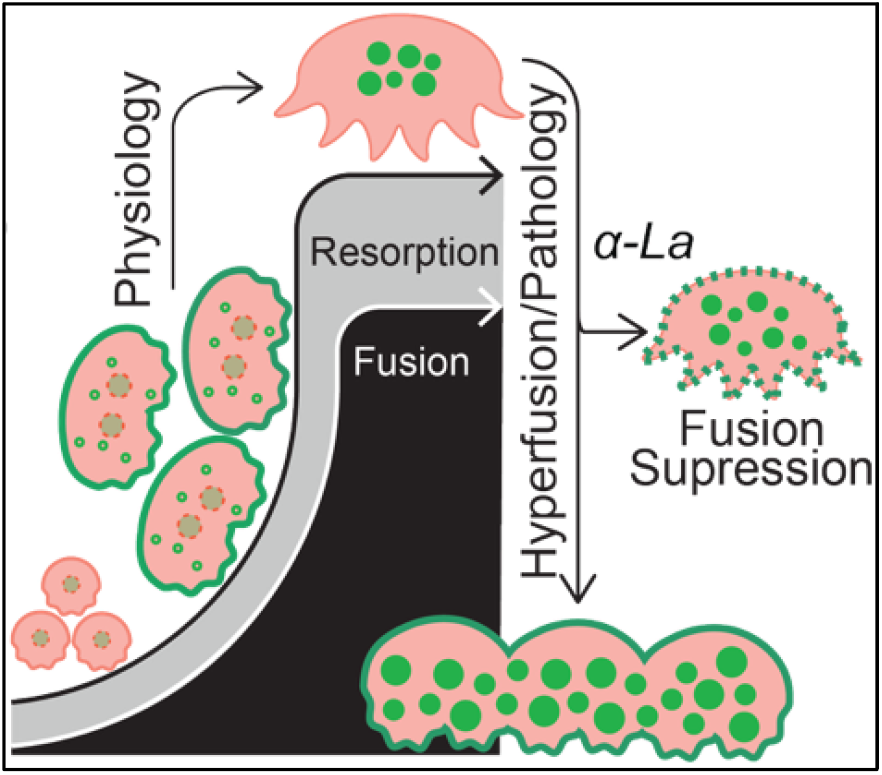
Osteopetrotic hyperfusion is facilitated by the retention of surface La. Cartoon depiction of La (green) surface trafficking – facilitating OC multinucleation – and removal upon OC reaching a mature size/functional range. In OC-rich ARO, La surface retention promotes hyperfusion and loss of resorptive capacity. La-targeted hyperfusion suppression rescues both OC size and resorptive function.

## Discussion

Perturbed OC multinucleation leads to alterations in skeletal metabolism and pathophysiology in a variety of skeletal diseases^10, 13-18^. Recent observations by Reuven *et al*. demonstrated that OC-rich ARO stemming from loss of SNX10, OSTM1, and CLC-7 function results in a similar hyperfusion phenotype leading to OCs that continuously multinucleate, never mature, and remain resorption incompetent^33^. Moreover, this recent report describes unregulated fusion between large, multinucleated OCs in response to SNX10, OSTM1, or CLC-7 loss, where each fusion event leads to substantially elevated osteopetrotic OC size compared with the typical fusion of multinucleated OCs with mononucleated cells^33^. Here, we report that hyperfusion in two genetic linkages of OC-rich ARO is underpinned by elevation of the multinucleation manager La at the surface of osteopetrotic OCs. Moreover, we demonstrate that suppressing elevated surface La rescues hyperfusion and mineral resorption competence. We view these *in vitro* data and our previous work assessing the efficacy of La inhibition in fibrous dysplasia of bone^22^ as proofs-of-principle supporting OC fusion as a viable mechanistic target in treating OC malformation and function in skeletal disease. In addition, since cell fusion is a relatively late stage in the formation of mature OCs, the effects of fusion-suppression on OC function and OC-to-osteoblast signaling can be more tunable when compared to treatments targeting pre-fusion stages of osteoclastogenesis.

Previous reports have demonstrated that OCs with loss of SNX10 function exhibit defects in endocytosis, and that CLC-7 and OSTM1 play important roles in the acidification of lysosomes^30, 31, 40^. Moreover, each of these proteins traffics to late endosomes/lysosomes and exhibits some co-localization with the others^33, 40^. Together, we agree with previous suggestions that OC-rich ARO may stem from a common defect in endo-lysomal trafficking. Perturbations in this endosomal sorting system following loss of *SNX10* or *OSTM1* may promote the retention of La (and probably other components of fusion machinery) at the surface of osteopetrotic OCs – leading to hyperfusion and the subsequent impairment of resorption competence. Alternatively, La and other OC fusion machinery may be degraded by the exocytosis of proteolytic enzymes that cleave away rather than internalize fusion proteins. In this alternative, loss of *SNX10* or *OSTM1* may impair the delivery of these degradative enzymes, leading to the impaired removal of surface La and the promotion of hyperfusion. Our data demonstrate that the hyperfusion resulting from this perturbed internalization of OC fusion machinery is not a parallel phenotype with impaired resorption, but rather, that hyperfusion contributes to resorption impairment and that resorption can be rescued by suppressing hyperfusion.

Our work has identified La as a highly specific, surface accessible target for modulating OC multinucleation and function. We propose that OC multinucleation represents a mechanistic target for modulating OC size and resorptive function in pathologic conditions. What if *instead of poisoning or ablating* OCs we could simply *tune their size and appetite* from pathological back into a healthy, physiological range in pathologies where OC size/function is elevated? This **“keep the clasts, tame their bite approach”** is poised to shift the current paradigm of skeletal therapies that ablate OCs to postpone bone loss, leading to side effects and severe discontinuation consequences in therapy. Our data suggests that represents a therapeutic target capable of moving this “keep the clasts” hypothesis into reality, with the goal of complementing current standards of care.

## Acknowledgements

We want to thank Dr. Kamran Melikov for many fruitful discussions and the National Institutes of Health Department of Transfusion Medicine for isolating the monocytes used in this study. This research in the L.V.C.’s, A.E.’s and B.G.’s laboratories was supported by Grant No 2021168 from the United States-Israel Binational Science Foundation (BSF). This research was supported in part by the Intramural Research Program of the National Institutes of Health (NIH). The contributions of the NIH author(s) are considered Works of the United States Government. The findings and conclusions presented in this paper are those of the author(s) and do not necessarily reflect the views of the NIH or the U.S. Department of Health and Human Services. The research in J.M.W.’s laboratory was supported by the *Eunice Kennedy Shriver* National Institute of Child Health and Human Development, National Institutes of Health Grant No R00-HD110609.

## SUPPLEMENTARY DATA

**Supplementary Figure 1.**
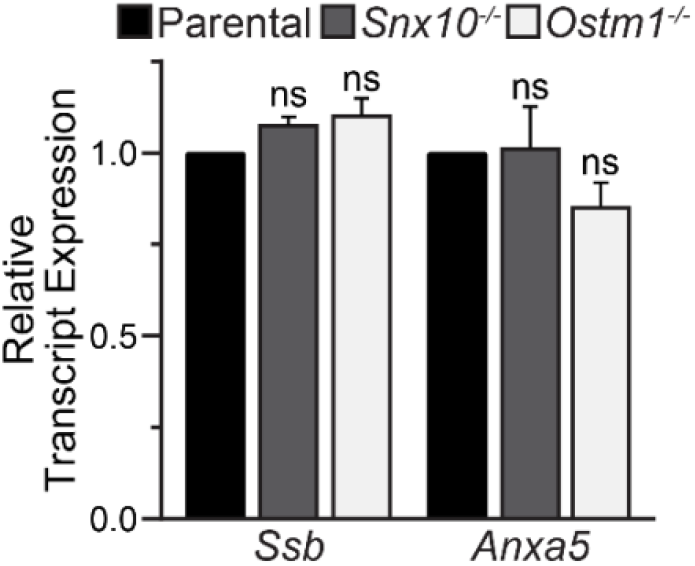
Quantification of the steady-state transcript levels of the osteoclast fusion regulators *Ssb* and *Anxa5* relative to *GAPDH*. n = 5. p = 0.06, 0.08, 0.95, and 0.21, respectively.

## METHODS

### Reagents

Murine and human M-CSF and RANKL were purchased from Cell Sciences (catalogue #s CRM146B; CRR100B; CRM735B and CRR101D). Mineral resorption assay kits were purchased from Cosmo Bio Co. (catalogue # CSR-BRA-24KIT). Molecular weight standards were purchased from Biodynamics (DM660).

### Cell Culture

RAW 264.7 cells (ATCC, Manassas, VA, # TIB-71) were maintained in Alpha MEM + nucleosides (Gibco # 12571063) supplemented with 10% FBS (Gibco # 26140079) to a maximum of 8 passages. Snx10-/-, Ostm1-/-, and their parental control clones were previously described^33^. RAW 264.7 cells were differentiated into osteoclasts in the presence of 100ng/ml murine RANKL for 5 days. Monocytes were elutriated from the peripheral blood of healthy donors were obtained through the Department of Transfusion Medicine at National Institutes of Health under protocol 99-CC-0168 approved by the National Institutes of Health Institutional Review Board. Research blood donors provided written informed consent and blood samples were de-identified prior to distribution, Clinical Trials Number: NCT00001846. We also used elutriated monocytes from healthy donors obtained through Elutriation Core Facility, University of Nebraska Medical Center, informed consent was obtained under an Institutional Review Board approved protocol for human subject research 0417-22-FB. Research blood donors provided informed consent and samples were de-identified prior to distribution. Monocytes were plated at 1×10^6^ per milliliter in Alpha MEM + nucleosides supplemented with 10% FBS, Penicillin-streptomycin-L-glutamine (Gibco # 10378016) and 100ng/ml human M-CSF. Media were replaced every 3^rd^ day for a total of 6 days. Following 6 days, adherent macrophage precursors were cultured in the same media + 100ng/ml human RANKL for up to 5 days.

### RNAi

siRNA were introduced into osteoclasts between days 2 and 3 of RANKL stimulation via Lipofectamine RNAiMAX according to the manufacturer’s instructions (ThermoFisher Scientific). Non-targeted (Cat#4390843), Snx10-targeted (Cat#4390771_ID:s211922), Ssb-targeted (Cat#4390771_ID:s74419), SNX10-targeted (Cat#4392420_ID:s26643), OSTM1 (Cat#4392420_ID:s26304) siRNA were introduced at a concentration of ∼15 ng/ml (Silencer Select, Ambion).

### Antibodies

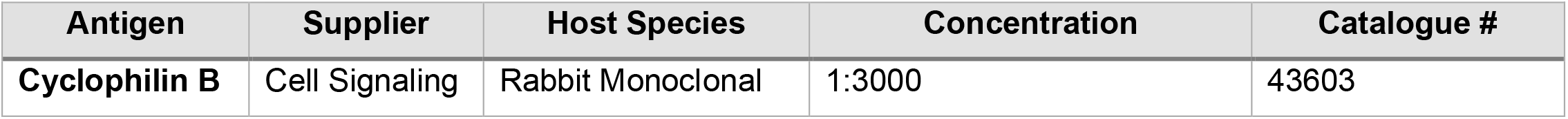
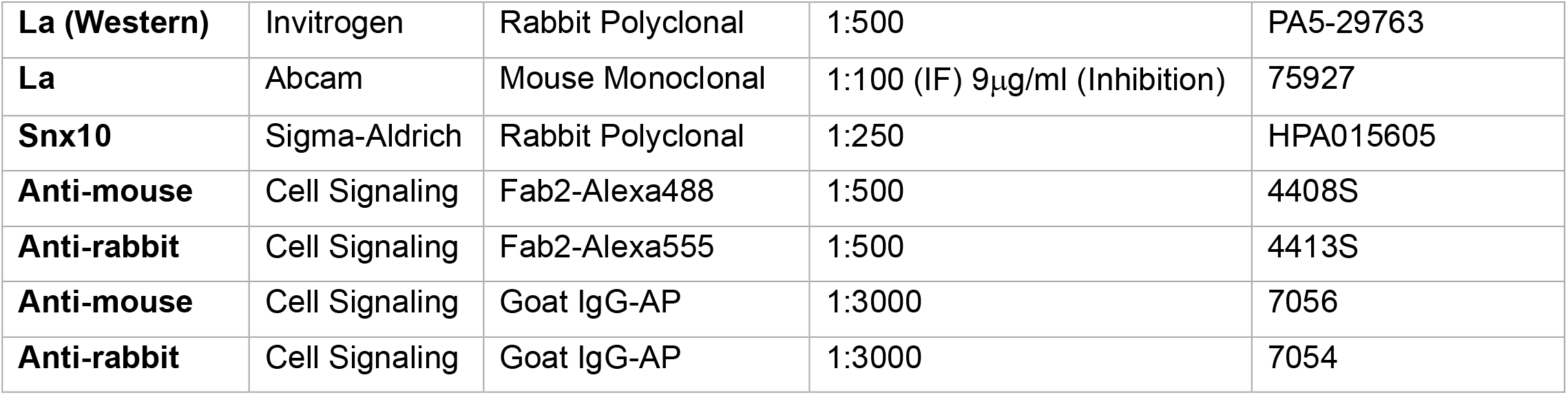

### Fluorescence Microscopy Imaging

For fluorescent imaging, cells were rapidly fixed with warm, freshly prepared 4% paraformaldehyde (Electron Microscopy Sciences Cat # 15710) in PBS. Cells were then washed with PBS and, in some cases, permeabilized via incubation with 0.1% Triton X100 in PBS. Some cells were then stained with phallodin-Alexa488 and Hoechst for quantifying cell multinucleation. For immunofluorescent imaging, cells are fixed and sometimes permeabilized as described above. Cells were then washed 3x with PBS and placed into PBS + 10% FBS for 10 min at room temperature to reduce non-specific binding. Cells were subsequently stained with primary antibodies for 1 hour at room temperature in PBS +10% FBS. Cells were then washed 5x in PBS and subsequently stained with secondary antibodies for 1 hour at room temperature in PBS +10% FBS. Cells were then washed 5x in PBS.

For multinucleation/fusion event quantification, 8 randomly selected fields of view were imaged using Alexa488, Hoechst and phase contrast compatible filter sets (BioTek) on a Lionheart FX microscope using a 10x/0.3 NA Plan Fluorite WD objective lens (BioTek) using Gen3.10 software (BioTek). The total number of nuclei in multinucleated osteoclasts with 3+ nuclei are divided by the total number of multinucleated osteoclasts with 3+ nuclei to obtain the average number of nuclei per osteoclast for each condition and then normalized to the control condition for the experiment. For immunofluorescent microscopy, images were captured on a Zeiss LSM 800, confocal microscope using a C-Apochromat 63x/1.2 water immersion objective lens.

All image data were evaluated using Fiji/ImageJ’s open-source image processing package v.2.1.0/153c.

### Mineral Resorption

Mineral resorption was evaluated utilizing Cosmo Bio USA resorption assay kits according to the manufacturer’s instructions. Briefly, Human, monocyte-derived osteoclasts were differentiated and treated with siRNA as described above using Alpha MEM without phenol red. Media were changed the morning after siRNA delivery (day 3-4post initial RANKL addition) and resorption conditioned media were collected at 4-5 days post initial RANKL addition, and fluorescence intensity within the media was evaluated as recommended by the manufacturer.

### Syncytin 1 Inhibitory Peptides

We used synthetic peptide inhibitor of Syn-1-mediated fusion (Syn-1 peptide, Ac-SGIVTEKVKEIRDRIQRRAEELRNTGPWGL-NH2) - described and characterized previously^37, 41^ - and, as a negative control (Syn-1 scr peptide), a peptide with the same amino acid composition but a scrambled sequence (Ac-GKWGLSRIRTELRNTEPVKEQVRAEIGDRI-NH2). Both peptides were used at 30μg/ml.

### Statistical Analysis

Statistical analyses were performed using Prism software (GraphPad Prism version 10.4.2). Unless stated in the legend, differences between groups were observed in each experiment, cells from the same source were paired across conditions, and statistical significance was assessed via Student’s t-test (in experiments comparing one control with one experimental condition) or ANOVA (in experiments comparing one control with multiple experimental conditions). Due to the inherent variability in the derivation of osteoclasts stemming from differing passages, donors, and/or precursor densities, we often analyzed statistical significance using a ratio paired test, where the raw values were logarithmically transformed and then assessed. The quantified results presented are all represent the mean ± the standard error of the mean. The p values for each statistical comparison are defined in the legends of each figure, however we graphically represented our statistical evaluations using the following symbols: ns, p = >0.05; * p = ≤0.05; **p = <0.01; ***p = <0.001; ****p = <0.0001. In each graph represented, n = the number of independent biological replicates.

## Notes

### Competing Interest Statement

The authors have declared no competing interest.

### Summary of Updates

For some reason one author name auto-populated in all caps. I simply corrected the capitalization of the name.

